# Grasshopper genome reveals long-term conservation of the X chromosome and temporal variation in X chromosome evolution

**DOI:** 10.1101/2022.09.08.507201

**Authors:** Xinghua Li, Judith E. Mank, Liping Ban

**Author notes:** To whom correspondence should be addressed (LB).

## Abstract

We present the first chromosome-level genome assembly of the grasshopper, *Locusta migratoria*, one of the largest insect genomes. We use coverage differences between females (XX) and males (X0) to identify the X chromosome gene content, and find that the X chromosome shows both complete dosage compensation in somatic tissues and an underrepresentation of testes-expressed genes. Remarkably, X-linked gene content from *L. migratoria* is highly conserved across four insect orders, namely Orthoptera, Hemiptera, Coleoptera and Diptera, and the 800 Mb grasshopper X chromosome is homologous to the fly ancestral X chromosome despite 400 million years of divergence, suggesting either repeated origin of sex chromosomes with highly similar gene content, or long-term conservation of the X chromosome. We use this broad conservation of the X chromosome to test for temporal dynamics to Fast-X evolution, and find evidence of a recent burst evolution for new X-linked genes in contrast to slow evolution of X-conserved genes. Additionally, our results reveal the X chromosome represents a hotspot for adaptive protein evolution related migration and the locust swarming phenotype. Overall, our results reveal a remarkable case of conservation and adaptation on the X chromosome.

## Introduction

Grasshoppers (order Orthoptera, suborder Caelifera) represent an important phylogenetic and developmental comparison to many insect model systems. The first grasshoppers likely arose 250 million years ago during the Triassic period (Mis of et al. 2014), and species within the group have since become some of the most prevalent herbivores on earth. The suborder, which contains more than 12,000 species, exhibits a worldwide distribution, with the greatest diversity in the tropics.

Grasshoppers normally possess XX/X0 sex chromosomes (Mao et al. 2020). X0 sex determination systems are thought to derive from XY systems with highly differentiated X and Y chromosomes in species where sex is determined based on X chromosome dose rather than Y-chromosome gene content (Furman et al. 2020). Because the Y chromosome is completely lost in X0 systems, they represent the ultimate example of sex chromosome heteromorphy. Extreme examples are often useful in revealing evolutionary patterns, however, despite their inherent utility for the study of sex chromosome, X0 sex chromosomes are relatively rare compared to XY systems (Bachtrog et al. 2014; The Tree of Sex Consortium 2014) and therefore their dynamics are not well understood. For example, although theory predicts that extreme heteromorphy will accelerate Fast-X evolution (Charlesworth et al. 1987) and the evolution of dosage compensation (Charlesworth 1996), empirical tests of this remain rare (Pal and Vicoso 2015).

Grasshoppers are also an excellent model organism for the study of phenotype plasticity. One of the most fascinating features within this clade is the phenomenon of locust swarming (Pener and Simpson 2009), the formation of dense migrating masses of grasshoppers that exhibit density-dependent phenotypic plasticity, known as locust phase polyphenism (Uvarov 1966; Perner 1983; Pener and Simpson 2009) which often cause extensive crop damage and food insecurity. Swarming locusts can migrate long distances, even between continents, and migratory locusts are broadly distributed throughout Africa, Asia, Europe, Australia, and nearby islands. Locust species belong to several different subfamilies of the family of Acrididae, and locust phase polyphenism presumably has evolved several times, by convergent, or partially convergent, evolution (Pener and Simpson 2009; Song et al. 2017).

Grasshoppers were an early genetic model, and Walter Sutton proposed the chromosome theory of heredity based in part on his work on grasshoppers at the start of the 20^th^ century (Crow and Crow 2002). Sutton’s success was partly attributed to the large chromosomes in grasshoppers, which result from extreme genome size, which in turn has hampered effective genome assembly and subsequent molecular studies. As a result, the two currently available grasshopper genome assemblies, *Locusta migratoria* (Wang et al. 2014) and *Schistocerca gregaria* (Verlinden et al. 2021) remain fragmented.

In this study, we combined the PacBio HiFi reads (Wenger et al. 2019) and Hi-C technology (Belton et al. 2012) to assemble the first high-quality chromosome-level genome of a grasshopper, the migratory locust, *Locusta migratoria*. Our genome assembly allows unprecedented insight into the role of extreme heteromorphism in sex chromosome evolution, and our results reveal surprising widespread conservation of the X chromosome gene content across broad swathes of the insect phylogeny as well as temporal dynamics to the rate of X chromosome evolution. We also combine our high-quality genome with extensive transcriptome data to identify positive selection for locust swarming phenotypes, revealing the underpinnings of this major form of phenotypic plasticity.

## Results

### Genome features

We used PacBio HiFi sequencing to generate genome sequences for a female (XX) migratory locust, and then used Hi-C reads to scaffold the contigs into a chromosome-level genome assembly comprising 12 chromosome-level scaffolds (Fig 1a). The final assembled genome size is 6.3 Gb with a contig N50 value of 52.8 Mb, the largest to date among published chromosome-level insect genome assemblies. To assess the completeness of our assembly, we performed BUSCO analyses against the insect orthologous groups and recovered a score of 96%, a major improvement on the previous migratory locust and desert locust assemblies (Fig 1b). Using our own and previously published RNA-seq datasets, we identified 26,636 protein coding genes with a total of 37,981 transcripts and 59,466 UTRs. Among the 26,636 genes, 19,481 were annotated by blastp against to the refseq arthropod proteins, including top-hits to *Zootermopsis nevadensis*.

**Fig 1.**
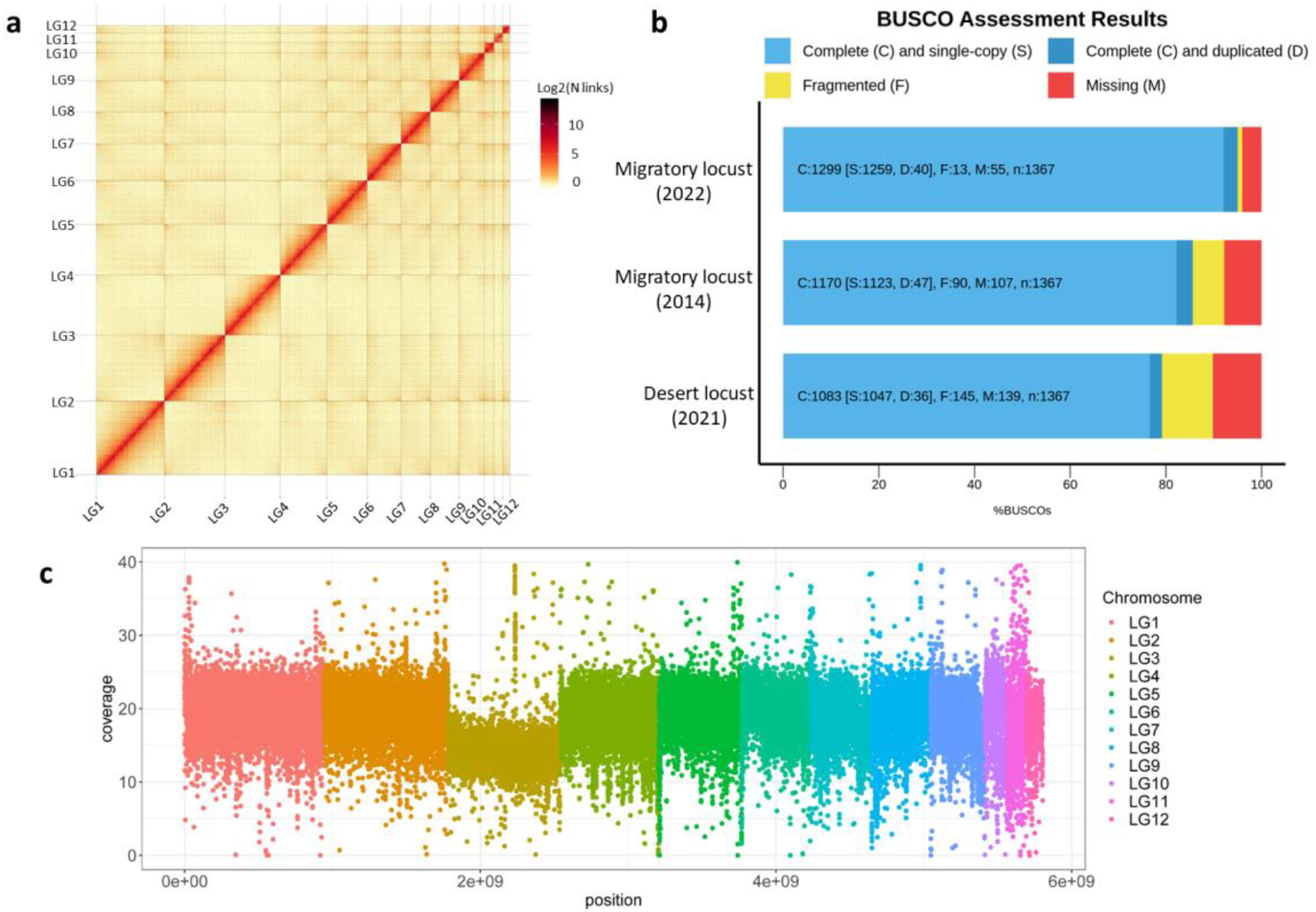
Chromosome-scale genome assembly. **a**, Hi-C contact map comprise 12 chromosome-level scaffolds; **b**, BUSCO assessment of our assembly, the previous migratory locust and desert locust assemblies; **c**, Male read depth along the genome in 100Kb windows.

The proliferation of repetitive elements is the main reason for the large size of the *L. migratoria* genome, and repetitive elements constituted 76.57% of the assembled genome, of which DNA transposons (19.87%) and LINE retrotransposons (28.13%) were the most abundant elements. The total repetitive content is much higher than previous reported (60%) (Wang et al. 2014), showing the advantage of the PacBio HiFi reads in assembly of high repetitive genomes. To investigate the genome quality, we also quantified the satellite DNA distribution along each chromosome (Fig S1). The most dominant satellites are LmiSat02A-176 and LmiSat27A-57. Surprisingly, we also identified several centromere and telomere specific satellites (namely LmiSat01A-185 and LmiSat07A-5-tel), suggesting that centromere and telomere repetitive elements have successfully integrated into some chromosomes, further demonstrating the high quality of our genome assembly.

### X chromosome identification and characteristics

To identify the X chromosome, we sequenced a male (X0) to an average of 30X coverage, mapping the Illumina reads to our genome and calculating read depth in 100Kb windows. Chromosome 3 has read depth nearly half of other chromosomes (Fig 1c), consistent with an X0 male karyotype and previous cytogenetic work (Cabrero et al. 2009).

We next compared features between the X chromosome and autosomes (Table S1; Fig S2). Compared to the autosomes, the X chromosome has lower gene density (T-test, P<0.001) and larger intron length (T-test, P<0.001). The population recombination rate (ρ) is lower on average across the X chromosome compared to all autosomes (T-test, P<0.001), except for LG4 and LG12 (T-test, P>0.05). The X chromosome also exhibits some differences in repetitive element distribution (Fig S2), with lower LINE transposon density (T-test, P<0.001) and higher DNA transposon density (T-test, P<0.001) compared to the autosomes. Interestingly, the Maverick transposon is significantly enriched on the X chromosome (T-test, P<0.001), where it is nearly twice as dense compared to the autosomes.

Next, we calculated the Kimura two parameter (K2P, (Kimura 1980)) distance of all transposons (Fig S3). The profile of X chromosome is similar to the small chromosomes, with a wave of Helitron proliferation in both chromosome classes.

### X-linked gene conservation across insect orders

We identified the conversation of *L. migratoria* X-linked gene content across four insect orders, Orthoptera, Hemiptera, Coleoptera and Diptera (Fig 2). In each comparison, the gene content shared on the X chromosome was greater than expected by chance based on the relative proportion of protein coding sites (chi-squared test, 1 d.f., P<0.05), suggesting either repeated origin of sex chromosomes with highly similar gene content, or long-term conservation of the X chromosome. Notably, the 800 Mb grasshopper X chromosome shares significant gene content to Muller element F in *D. melanogaster* (the ancestral fly X chromosome, (Vicoso and Bachtrog 2013)) (chi-squared test, 1 d.f., P=3.84×10^-31^) despite 400 million years of divergence. Through functional enrichment analysis, we show that these conserved X-linked genes include GO terms such as learning and memory, neuron recognition and growth hormone synthesis (Fig S4).

**Fig 2.**
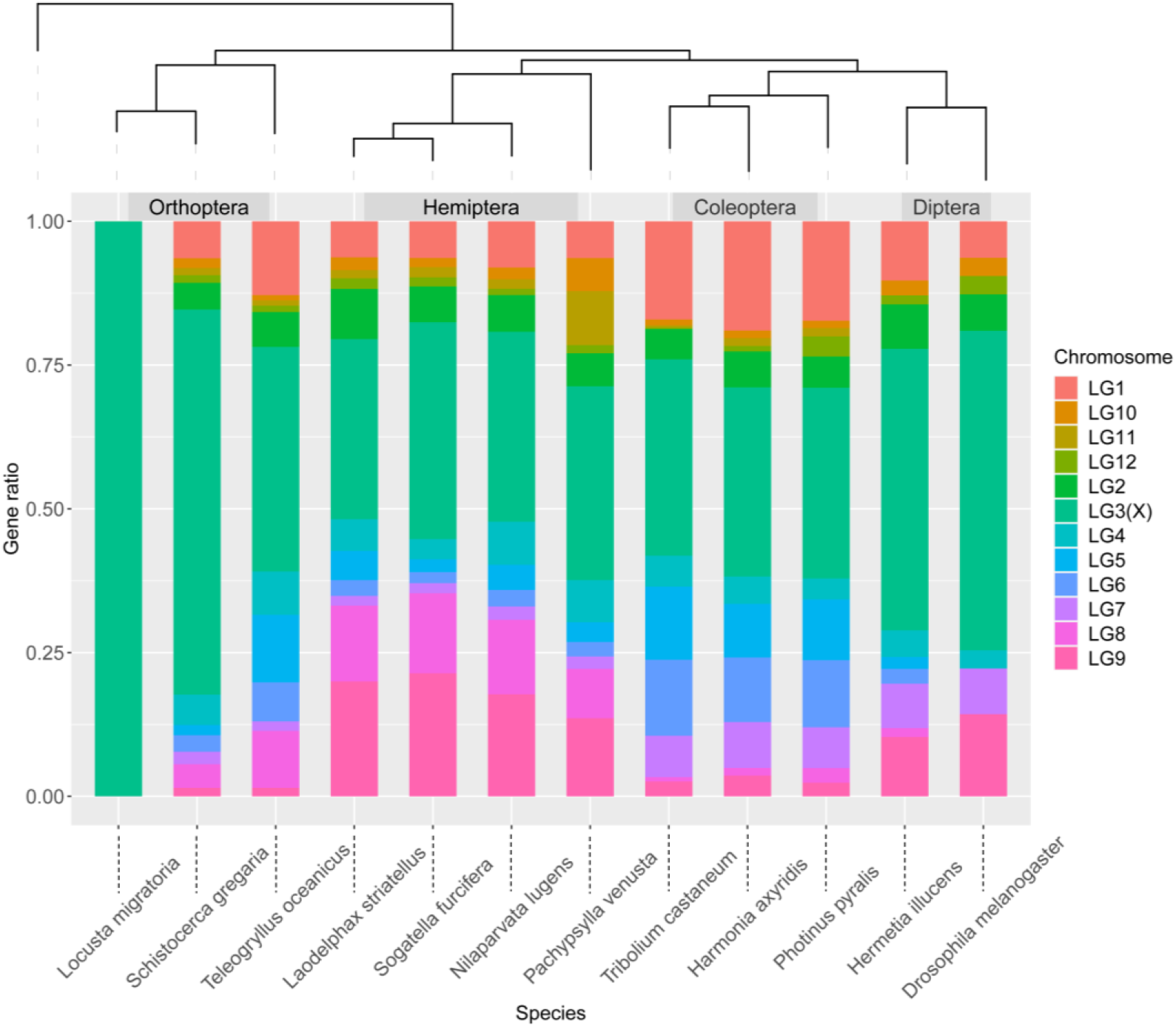
X-linked gene content conservation across four insect orders. The proportion of X-linked genes of each species with the genomic location of *L. migratoria* homologs identified by reciprocal best hit. Genome comparisons include *Schistocerca gregaria*(Verlinden et al. 2020), *Teleogryllus oceanicus*(Pascoal et al. 2020), *Laodelphax striatellus*(Zhu et al. 2017), *Sogatella furcifera*(Wang et al. 2017), *Nilaparvata lugens*(Ye et al. 2021), *Pachypsylla venusta*(Li et al. 2020), *Tribolium castaneum*(Richards et al. 2008), *Harmonia axyridis*(M. Chen et al. 2021), *Photinus pyralis*(Fallon et al. 2018), *Hermetia illucens*(Generalovic et al. 2021) and *Drosophila melanogaster*(Celniker et al. 2002).

### Variation in the tempo of Fast-X Evolution

We classified genes in *L. migratoria* into five, partially overlapping, categories: X-conserved genes (Fig S4) are X-linked in at least eight species from Fig 2; X-Lmig are X-linked only in *L. migratoria* and autosomal in all other species from Fig 2; X-X genes are X-linked in *L. migratoria* and only one other species; A-A genes are autosomal in *L. migratoria* and all other species; A-X genes are autosomal in *L. migratoria* and X-linked in other species. For each of these categories, we calculated average d_N_/d_S_ (Fig 3, Table S2), comparing each category to X-Lmig via bootstrapping (1000 replicates).

**Fig 3.**
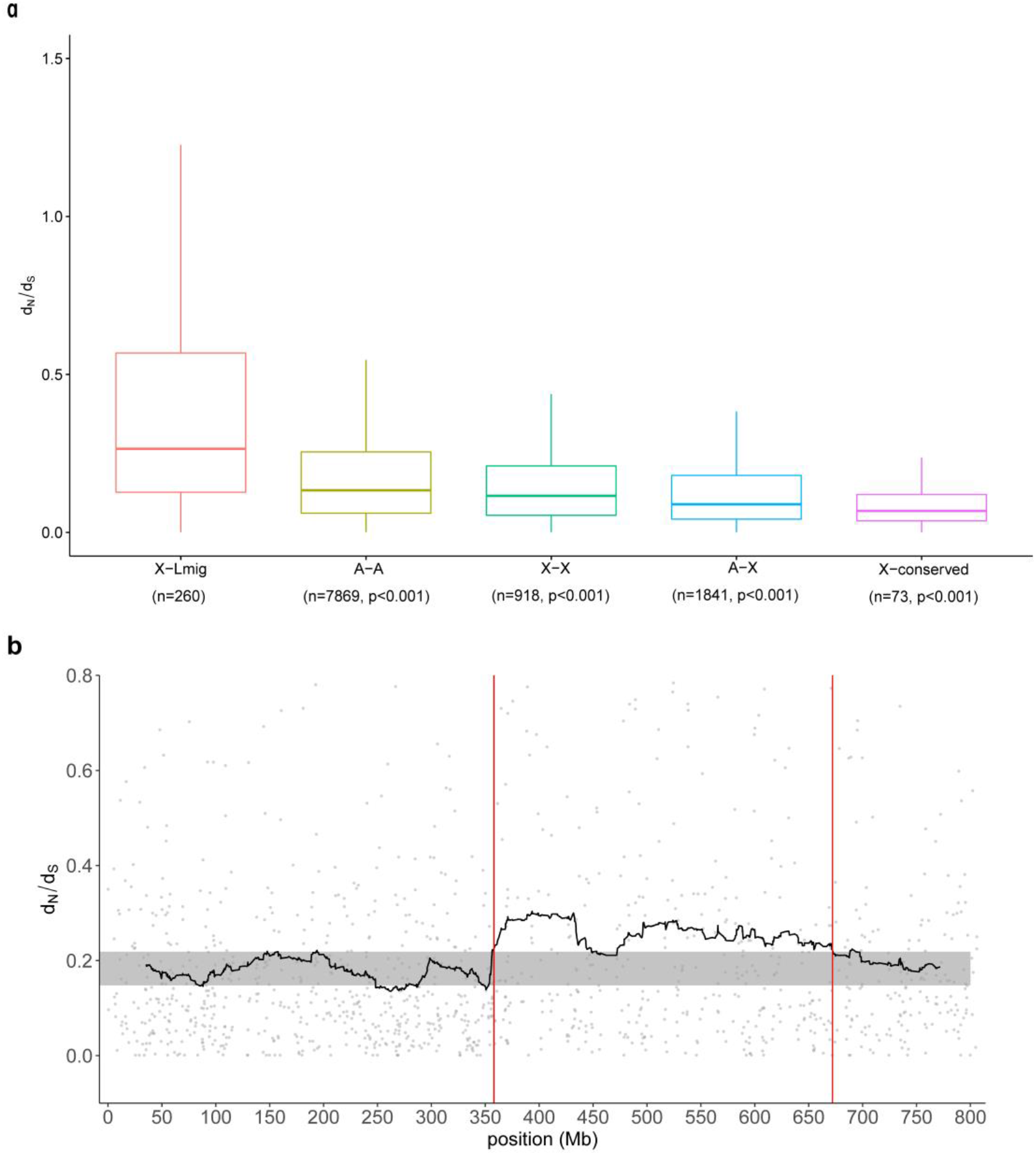
Gene evolution rate between X chromosome and autosomes. **a,** Boxplot of d_N_/d_S_ values from different categories; **b**, The moving average values of d_N_/d_S_ along the X chromosome.

We observe elevated rates of evolution in X-Lmig genes compared to all other categories of genes, consistent with Fast-X evolution (Fig 3, Table S2). Notably, we did not observe elevated d_N_/d_S_ for X-conserved or X-X genes, suggesting that Fast-X primarily results from genes specific to the *L. migratoria* X chromosome. Importantly, X-conserved genes showed significantly slower rates of average evolution compared to A-A genes (P=0.002), suggesting both Fast-X and Slow-X in the same species depending on the age of X-linkage. Fast-X and Slow-X are mainly due to differences in dN values (Table S2). To validate this, we performed the same analysis in true bugs (Hemiptera) and recovered similar results (Fig S5).

We next analyzed d_N_/d_S_ patterns across the X chromosome (Fig 3b), recovering a region (360MB ~ 670Mb) of elevated d_N_/d_S_ compared to both the autosomes (P<0.0001 based on 10,000 bootstraps) and the remainder of the X chromosome (P<0.0001 based on 10000 bootstrap replicates) level. This suggests that Fast-X might be explained by regional variation along the X chromosome.

### X Chromosome Dosage Compensation

We next tested for the presence of complete dosage compensation in *L*. *migratoria* (Fig 4). In female (XX) grasshoppers, X-linked genes showed higher expression levels than autosomal genes in both somatic and gonad tissues. In male (X0) grasshoppers, X-linked gene expression is higher than or equal to autosomal genes in somatic tissues, but significantly lower in testis (P<0.001). The overall male X-linked expression was equal to female X-linked expression in somatic tissues, but significantly lower in testis (P<0.001) consistent with complete X chromosome dosage compensation in somatic cells.

**Fig 4.**
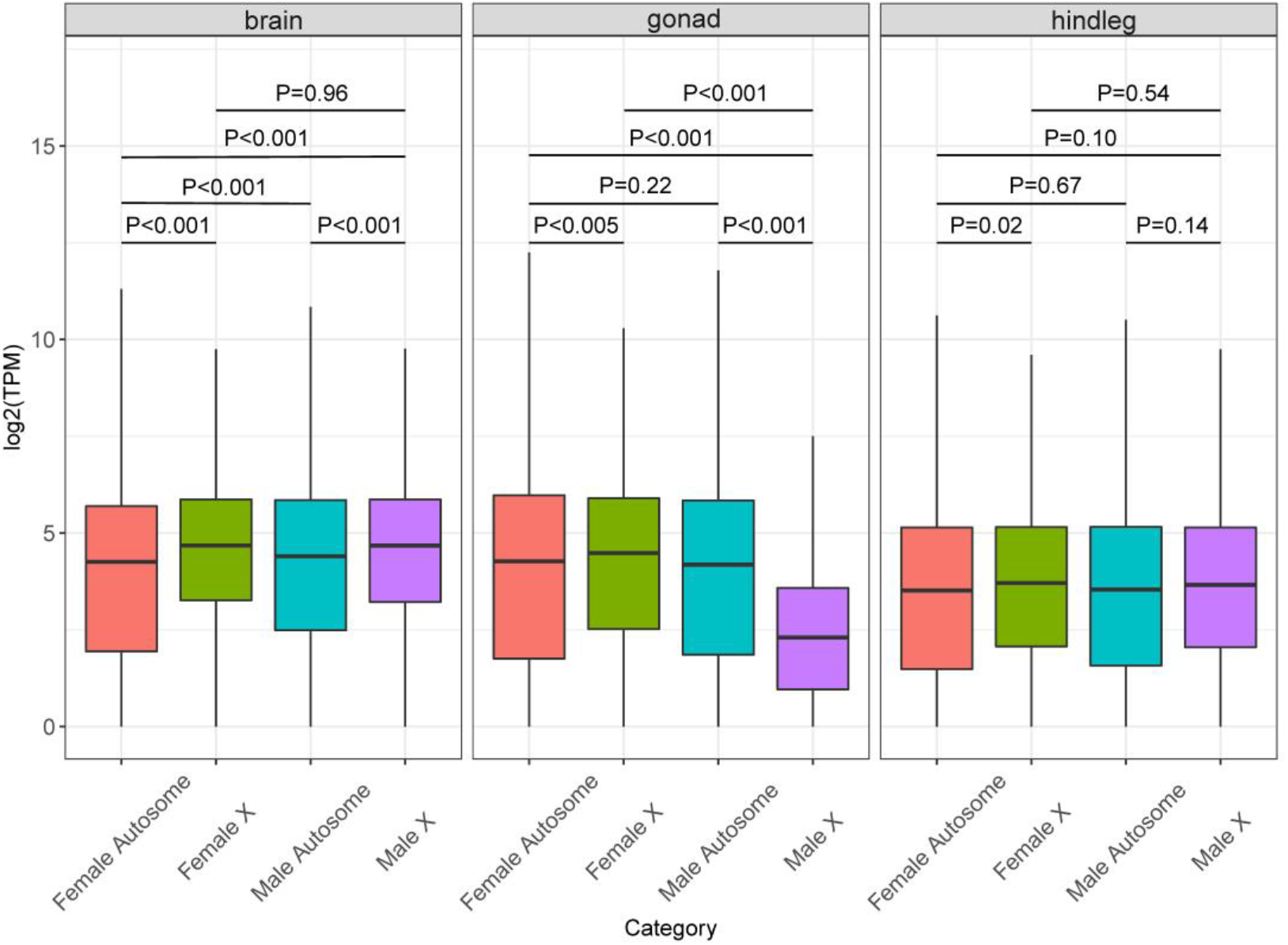
Dosage compensation in L. migratoria. P values were calculated based on 1,000 bootstrap replicates.

### Positive selection on the X chromosome

Based on our high-quality genome annotation, we carried out a phylogenetic analysis of the swarming locust phenotype in *L. migratoria* in comparison to three non-swarming species with transcriptome assemblies, namely *Oedaleus asiaticus*(Qin et al. 2017), *Gomphocerus sibiricus*(Shah et al. 2019) and *Xenocatantops brachycerus*(Zhao et al. 2018). After analyzing the migratory locust and three non-swarming grasshopper protein-coding sequences by PAML with an unrooted tree, (*Xenocatantops brachycerus*,(*Oedaleus asiaticus, Locusta migratoria* #1), *Gomphocerus sibiricus*), we detected 440 genes under positive selection. Of these 67 (15 %) are located on the X chromosome, representing a significant enrichment based on total gene content of the X (Chi square test, 1.df, P=0.009). Functional enrichment analysis of positively selected genes reveals GO terms including rRNA processing, cell cycle phase transition, muscle contraction, myosin filament assembly, olfactory transduction etc. (Fig S6).

## Discussion

### Hi-C and long read sequencing resolve a large complex insect genome into chromosomes

We used a combination of long-read DNA and Hi-C sequencing to successfully resolve and assemble an unusually large and highly repetitive insect genome. To date, this is the largest insect genome, and one of the largest arthropod genomes, assembled to chromosome scale. This is remarkable because the assembly of relatively large and highly repetitive insect genomes into highly contiguous chromosomes was until very recently unattainable, largely due to the difficulties presented by high amounts of repetitive content. Indeed, the unusually large size of the grasshopper genome is primarily due to the high proportion of repetitive content, corresponding to 76.57% of the genome. Using our new high-quality genome assembly of *Locusta migratoria*, we investigated X chromosome dynamics and adaptive evolution associated with the locust swarming phenotype.

### Surprising conservation of X chromosome gene content

We observe high conservation of X chromosome gene content across Insecta (Fig. 2). This is surprising, particularly given observations of turnover in sex chromosomes within the Diptera from the ancestral Dipteran X chromosome, the dot chromosome (Vicoso and Bachtrog 2015). However, conservation of X gene content has been observed in a limited number of Insecta orders (Chauhan et al. 2021), and our work illustrates a broader conservation across the Class. The extended evolutionary distance precludes a meaningful synteny analysis, and so it remains unclear whether this conservation in gene content reflects conservation of the sex chromosome itself, repeated origin of X chromosomes from the same underlying syntenic regions, or repeated movement of the same gene content to the sex chromosomes.

### Fast-X and Slow-X evolution

The X chromosome has several properties that distinguish it from the autosomes (Vicoso and Charlesworth 2006; Meisel and Connallon 2013) and that have the potential to influence the rate and pattern of evolution of X-linked genes (Charlesworth et al. 1987). Because males have only one copy of the X chromosome and therefore only one copy of X-linked genes, recessive mutations on the X chromosome are directly exposed to selection in males (Charlesworth et al. 1987). This can lead either to rapid fixation of recessive beneficial variation (Fast-X) or more efficient purging of recessive deleterious mutations (Slow-X) (Xu et al. 2012).

We observe Fast-X on genes that are X-linked only in *L. migratoria* and Slow-X for genes that are conserved on the X chromosome across insects (Fig 3a). This may suggest that the pool of adaptive recessive variation is quickly depleted following X-linkage, resulting in a limited burst of Fast-X. Over time, this dynamic appears to shift such that recessive deleterious variation is purged more effectively on the X, resulting in Slow-X over greater evolutionary distances. Interaction of Fast-X and Slow-X has previously been observed over far shorter timescales (Xu et al. 2012), however the extraordinary conservation of X chromosome coding content that we observe here across Insecta makes it possible to discern temporal dynamics in X chromosome evolution across extreme timespans. The temporal dynamics of X evolution that we observe, in addition to the fact that Fast-X in *L. migratoria* is largely confined to a restricted region (360MB ~ 670Mb, Fig 3B), suggests that this Fast-X region may represent a recent addition to the X chromosome.

Fast-X and Slow-X evolution is expected to be exacerbated in species with complete sex chromosome dosage compensation (Vicoso and Charlesworth 2009; Mank et al. 2010), which we observe for somatic tissues in *L. migratoria* (Fig 4), as well as the extreme heterogamety represented in X0 sex chromosome systems (Darolti et al. 2021). Fast-X, accelerated by dosage compensation and extreme heterogamety may in turn increase the role of the X chromosome in adaptation and speciation relative to its size and coding content, termed the Large-X effect (Lasne et al. 2017). Indeed, positively selected genes associated with the locust swarming phenotype are significantly enriched in X chromosome.

## Materials and Methods

### Library construction and sequencing

For PacBio sequencing, genomic DNA of a female migratory locust was isolated and sheared to an average size of 20 kb using a g-TUBE device (Covaris, Woburn, MA, USA). The sheared DNA was purified and end-repaired using polishing enzymes, followed by blunt end ligation and exonuclease treatment to create a SMRTbell template according to the PacBio 20-kb template preparation protocol. A BluePippin device (Sage Science, Beverly, USA) was used to size-select the SMRTbell template and enrich large (>10 kb) fragments. SMRTbell libraries were sequenced on a PacBio Sequel II system and consensus reads (HiFi reads) were generated using ccs software (https://github.com/pacificbiosciences/unanimity).

For Hi-C sequencing, Hi-C libraries were prepared from a male migratory locust at BioMarker Technologies Company (Beijing, China). Briefly, sample was collected and spun down, and the cell pellet was resuspended and fixed in formaldehyde solution. DNA was isolated and the fixed chromatin was digested with the restriction enzyme DpnII overnight. The cohesive ends were labeled with Biotin-14-DCTP using Klenow enzyme and then religated with T4 DNA ligation enzyme. Subsequent DNA was sheared by sonication to a mean size of 350 bp. Hi-C libraries were generated using NEBNext Ultra enzymes and Illumina-compatible adaptors. Biotin-containing fragments were isolated using streptavidin beads. All libraries were quantified by Qubit2.0, and insert size was checked using an Agilent 2100 and then quantified by quantitative polymerase chain reaction (PCR). Hi-C sequencing was performed by Illumina HiSeq 2500 platform, using paired-end of 150-bp reads.

To assist gene prediction and dosage compensation analysis, 24 RNA-sequencing (RNA-seq) libraries were generated from brain, hindleg and gonads with 4 biological replicates for each sex. Total RNA was extracted from each tissue using a TRIzol kit (Life Technologies, Carlsbad, USA). The mRNA fractions were isolated from the total RNA extracts with the MicroPoly (A) Purist kit (Ambion, TX, USA). cDNA libraries were prepared for each tissue with the RNA-seq Library kit (Gnomegen, San Diego, CA, USA) following the manufacturer’s instructions. Each paired-end cDNA library was sequenced with a read length of 150 bp using the Illumina HiSeq 2500 sequencing platform. All sequencing was performed by Biomarker Technologies Company (Beijing, China).

### Genome assembly

The PacBio long (~12 kb) and highly accurate (>99%) HiFi reads were assembled to a contig-level assembly using Hifiasm (Cheng et al. 2021). The Hi-C data were mapped to Hifiasm contigs with BWA (version 0.7.17-r1188). Uniquely mapped data were used for chromosome-level scaffolding. HiC-Pro (version 2.8.1) was used for duplicate removal and quality controls, and the filtered Hi-C data were then used to correct misjoins as well as to order and orient contigs. Preassembly was performed for contig correction by splitting contigs into segments with an average length of 300 kb, and then the segments were preassembled with Hi-C data. Misassembled points were defined and broken when split segments could not be placed to the original position. Then, the corrected contigs were assembled using LACHESIS with parameters CLUSTER_MIN_RE_SITES = 225, CLUSTER_MAX_LINK_ DENSITY = 2; ORDER_MIN_N_RES_IN_TRUN = 105; ORDER_ MIN_N_RES_IN_SHREDS = 105 with Hi-C valid pairs. Gaps between ordered contigs were filled with 100 “N”s.

To evaluate the quality of the genome assembly, we performed BUSCO (version v5.4.2) analyses using 1,367 core conserved insect genes on the old assembly (Wang et al. 2014), the recent desert locust assembly (Verlinden et al. 2020) and our assembly.

### Repeat annotation and gene prediction

De novo identification of repeats was performed by the RepeatModeler under default parameters. We also recovered 107 satellite DNA sequences belonging to 62 families in *L. migratoria* (Ruiz-Ruano et al. 2016). Using the ab initio repeat library and satellite DNA library, we estimated the repeat content of the assembled genome using RepeatMasker. Ab initio gene prediction was performed using Augustus. GenomeThreader, implemented in BRAKER, was run for homology-based prediction using protein sequences of *Drosophila melanogaster, Anopheles gambiae, Tribolium castaneum, Apis mellifera, Bombyx mori, Acyrthosiphon pisum* and *Zootermopsis nevadensis*. Publicly available NCBI transcriptome data and our own transcriptome data were aligned by HISAT2 and assembled with stringtie, and then coding regions were identified with TransDecoder. Finally, EVidenceModeler (EVM) was used to integrate the prediction results obtained with the above three methods. PASA (version v2.4.1) was run for gene structure annotation.

### X chromosome identification via coverage in males

To identify the X chromosome, a male migratory locust was sequenced to nearly 30X coverage. The Illumina reads were aligned to our genome assembly with BWA (version 0.7.17-r1188) and samtools was used to remove PCR duplicates. Mosdepth (Pedersen and Quinlan 2018) was used to calculate read coverage along the genome (parameters: -t 3 -n --fast-mode --by 500000).

### Gene density, GC content, nucleotide diversity

Gene density of each chromosome was calculated as the number of genes divided by chromosome length. GC content along chromosomes was calculated within 50 kb sliding windows. VCFtools (v0.1.13) was used to determine nucleotide diversity within 500 kb sliding windows.

### Recombination rate estimation

To explore the recombination across the locust genome, we estimated the population recombination rate (ρ) using FastEPRR (Gao et al. 2016). First, five female grasshopper resequencing data was download from NCBI Bioproject PRJNA433455. bcftools mpileup was used to call SNPs. Then Beagle (version 5.0) was used to phase the SNPs, and phased data were then input into the FastEPRR_VCF_step1 function in FastEPRR to scan each 10 and 50 Kb window (with parameters inSNPThreshold = 30 and qualThreshold = 20). Next, FastEPRR_VCF_step2 was used to estimate the recombination rate for each window. Finally, we applied FastEPRR_VCF_step3 to merge the files generated by step 2 for each chromosome.

### K2P analysis

RepeatMasker was used to construct the TE expansion history in the migratory locust genome by first recalculating the divergence of the identified TE copies in the genome with the corresponding consensus sequence in the TE library using Kimura distance and then estimating the percentage of TEs in the genome at different divergence levels.

### Gene content in insect orders

Insect genomes with assembled sex chromosomes were retrieved from InSexBase (X. i Chen et al. 2021). The proportion of X-linked genes of each species with their *L. migratoria* homologs were identified by reciprocal best blast hit, including *Schistocerca gregaria*, *Teleogryllus oceanicus*, *Laodelphax striatellus*, *Sogatella furcifera*, *Nilaparvata lugens*, *Pachypsylla venusta*, *Tribolium castaneum*, *Harmonia axyridis*, *Photinus pyralis*, *Hermetia illucens* and *Drosophila melanogaster*.

### Functional enrichment of genes

Gene functions the Gene Ontology (GO) annotations were retrieved with eggNOG-mapper (Cantalapiedra et al. 2021). Because *L*. *migratoria* is not a model organism, a local OrgDb database was constructed based on eggNOG-mapper results. The functional enrichment was then determined using clusterProfiler (Yu et al. 2012).

### Fast-X analysis

The evolution rate of genes was calculated by comparing the grasshopper *Oedaleus asiaticus*, which belongs to the same subfamily, *Oedipodinae*, as *L. migratoria*. Transcriptomic data of this species were downloaded from the NCBI SRA database (SRR IDs SRR2051024, SRR3372608, SRR3372609, and SRR3372610). Trinity was used to assemble a transcriptome representing a non-redundant gene set of this species. The reciprocal best blast hit pairs were used to Identify orthogroups. KaKs_Calculator (v2.0, https://sourceforge.net/projects/kakscalculator2/) was used to calculate d_N_/d_S_ values. Orthologous genes with d_N_/d_S_ > 2 were removed. The statistical tests between different gene categories were performed in R 4.1.1.

### Dosage compensation analysis

RNA-seq reads from heads, hindlegs and gonads of four females and four males were trimmed for adapter and low-quality bases (Q < 20) using fastp (Chen et al. 2018). Next, the RNA-seq reads were mapped to the genome using HISAT2 (Kim et al. 2019). Abundance estimation was performed with FeatureCounts (Liao et al. 2014). The raw counts were normalized by TPM methods. Genes with low expression support (sum of normalized read count of all samples < 1) were removed from downstream analysis. Dosage compensation was assessed by comparing average expression between female autosomal and X genes, male autosomal and male X genes, female autosomal and male autosomal genes, and between female X and male X genes.

### Positive Selection

To detect the evolutionary feature of grasshoppers, we used our genome and 3 non-swarming grasshoppers (*Oedaleus asiaticus*, *Gomphocerus sibiricus* and *Xenocatantops brachycerus)* to identify positive selection. Trinity was used to assemble transcriptomes representing a non-redundant gene set of these species. All the orthologues from the results of the reciprocal best hit (RBH) method were used to test for positive selection. One-to-one orthologues for the 4 species were aligned by MAFFT and gaps were removed by Gblocks v0.91b. The species tree generated by RAxML was used as the input tree for positive selection. The branch-sites model in PAML was used to look for positive selection. Multiple testing was corrected by Benjamini and Hochberg’s False Discovery Rate.

## Authors’ contributions

LB conceived and designed the research. XL performed the experiments, analyzed and interpreted the data, and wrote the manuscript. JM analyzed the data and wrote the manuscript. All authors read and approved the final manuscript.

## Acknowledgments

This work was supported by the earmarked fund for the National Key Research and Development Program of China and Beijing Agriculture Innovation Consortium (BAIC02-2022).

