## Supplementary files for "Grasshopper genome reveals long-term conservation of the X chromosome and temporal variation in X chromosome evolution"

,

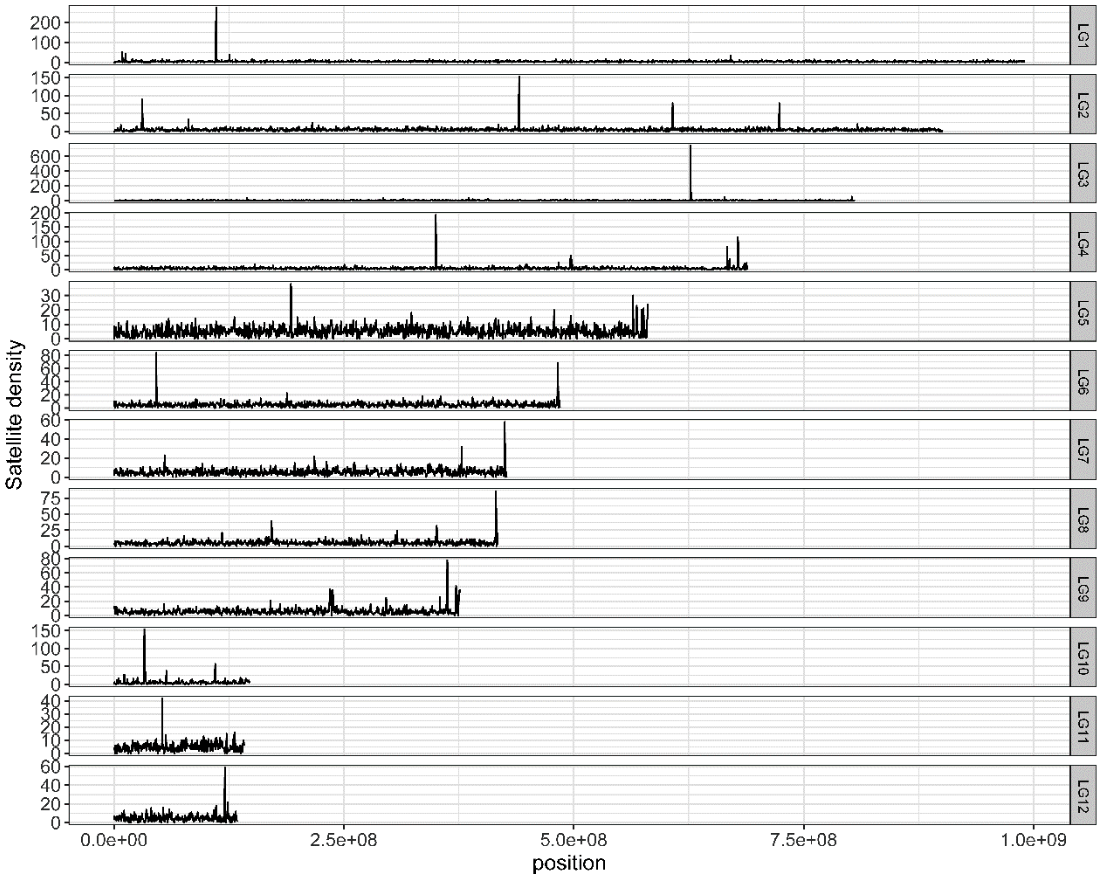


**Fig S1. Satellite distribution along chromosomes in the migratory locust genome.**

**a**

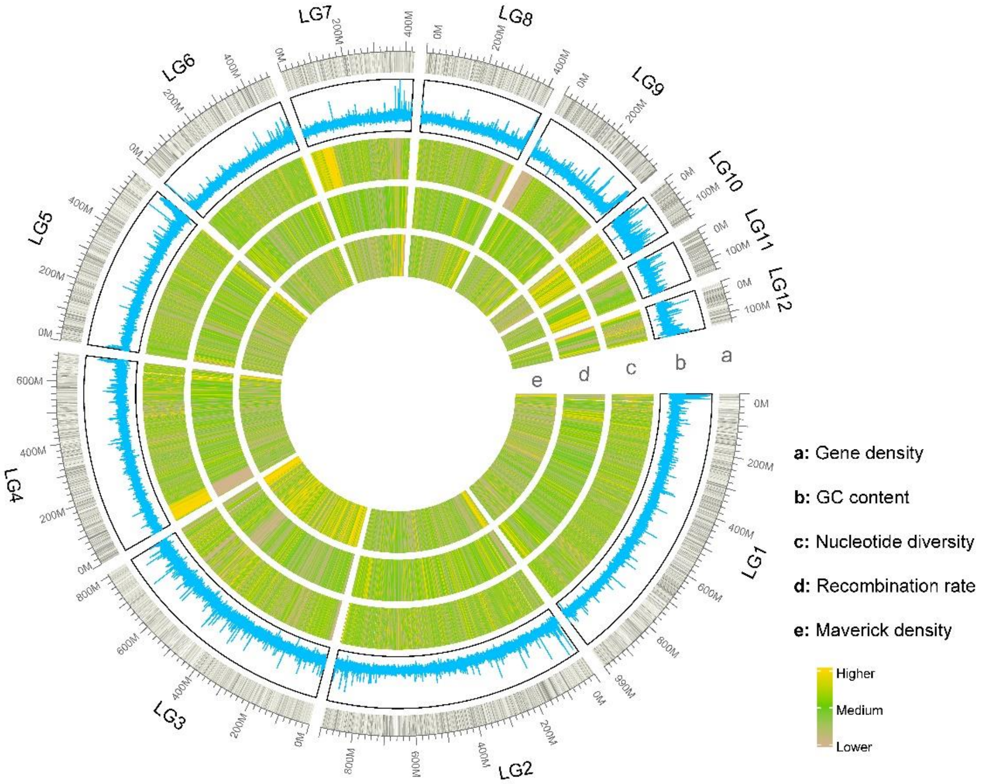


**b**

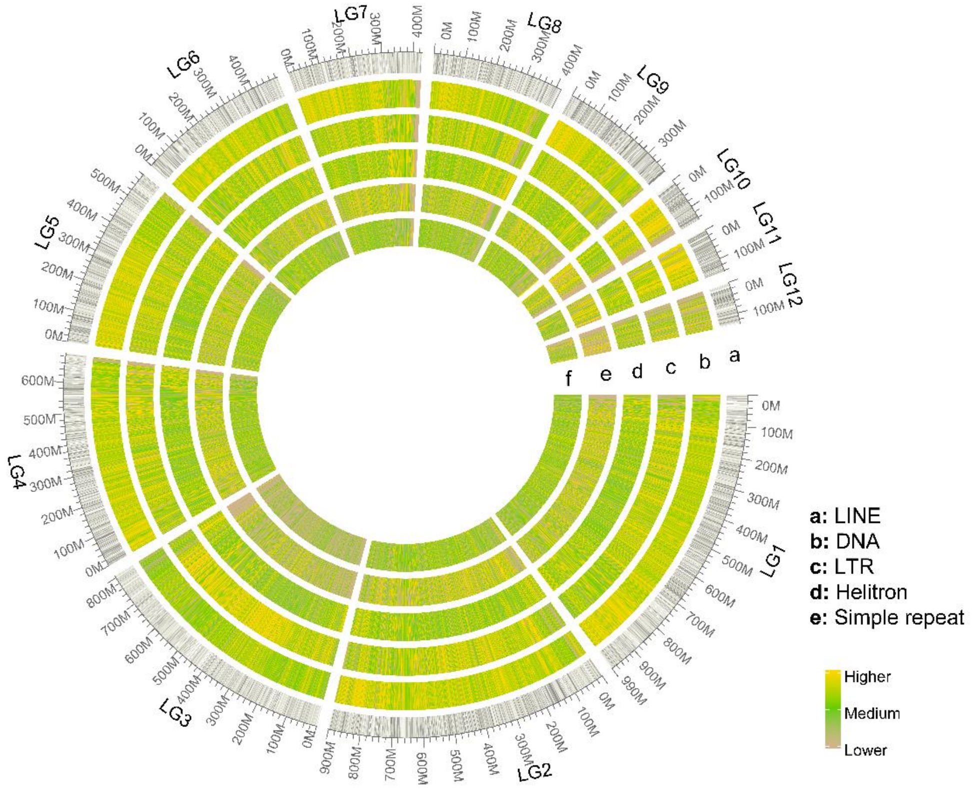


**Fig S2. Genomic features along migratory locust chromosomes. a**, gene density, GC content, nucleotide diversity, recombination rate; **b**, TE profile including LINE, DNA, LTR, Helitron transposons and simple repeat.

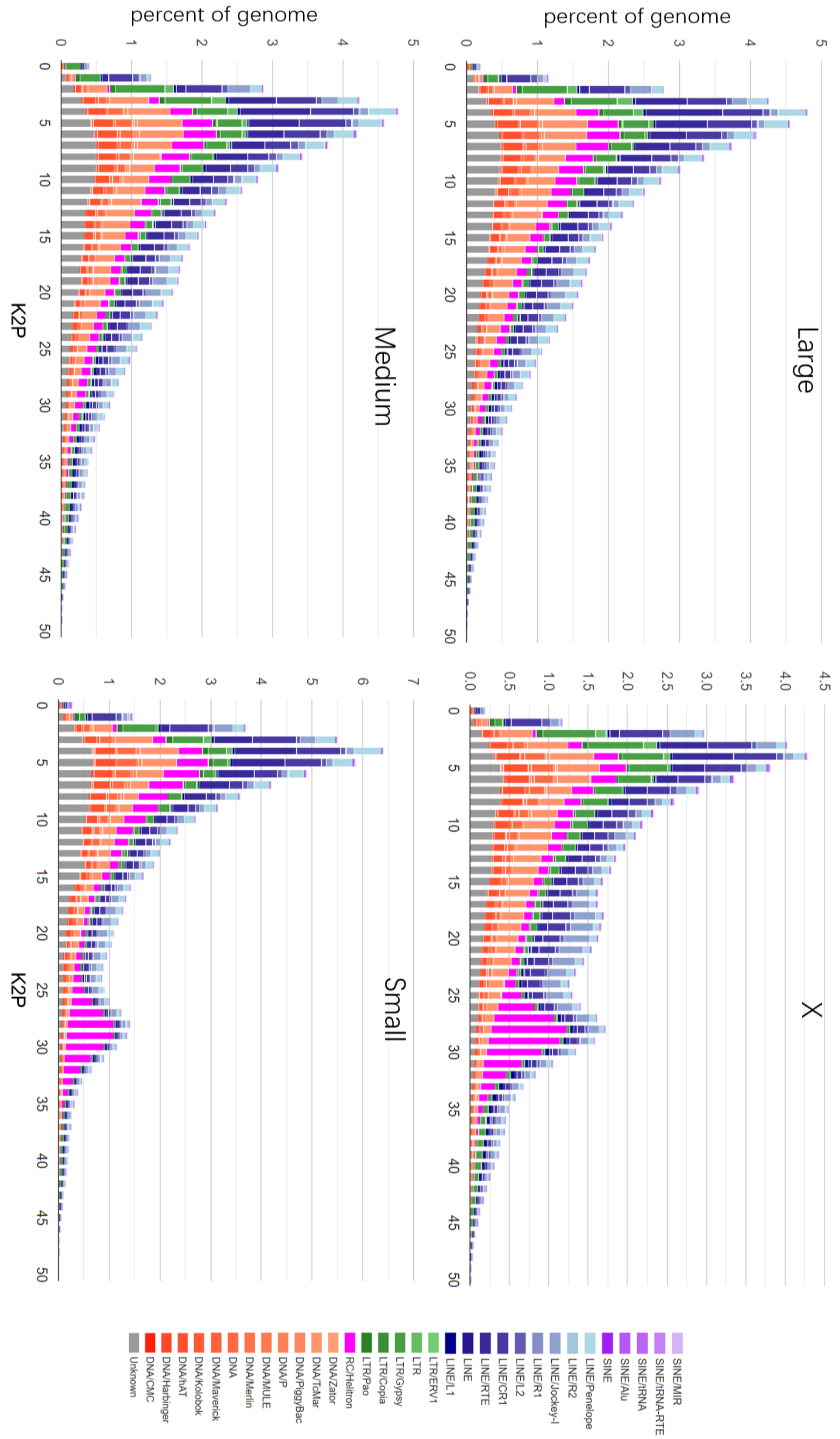


**Fig S3. TE percentage and K2P profile.** According to the size, chromosomes were classified into four categories: Large, Medium, Small and X.

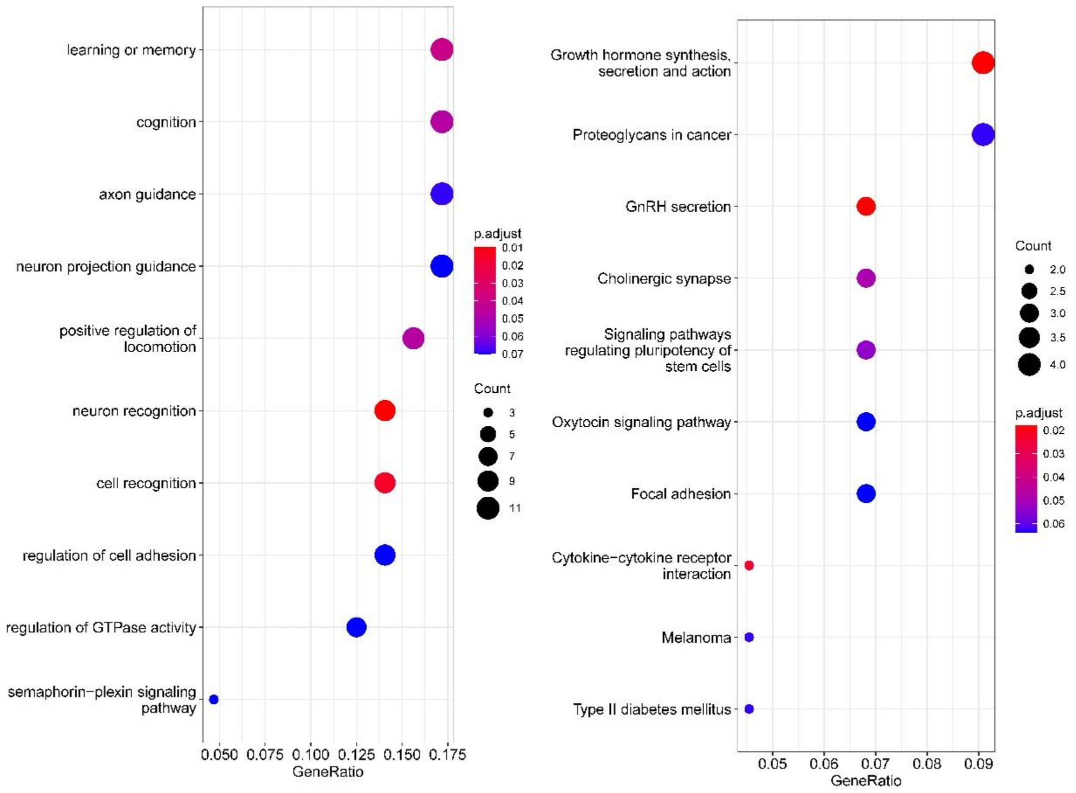


**Fig S4. Functional enrichment of conservative X-linked genes.** Left is GO terms, right is KEGG terms.

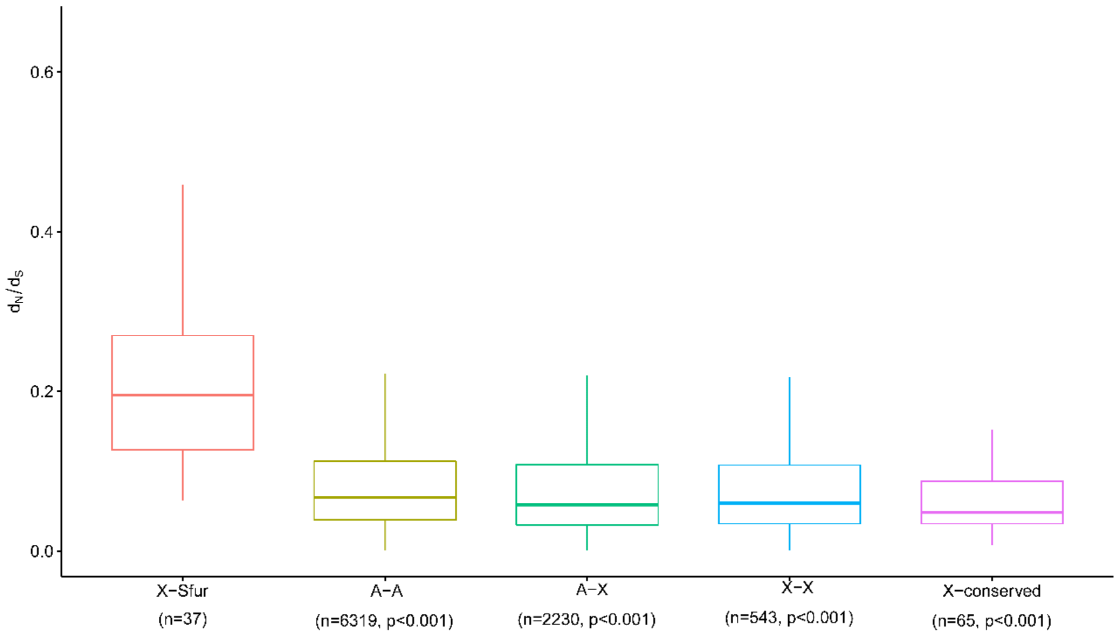


**Fig S5. dN/dS analysis in true bugs.** P values were calculated between X-Sfur and others by 1000 replicates bootstrapping.

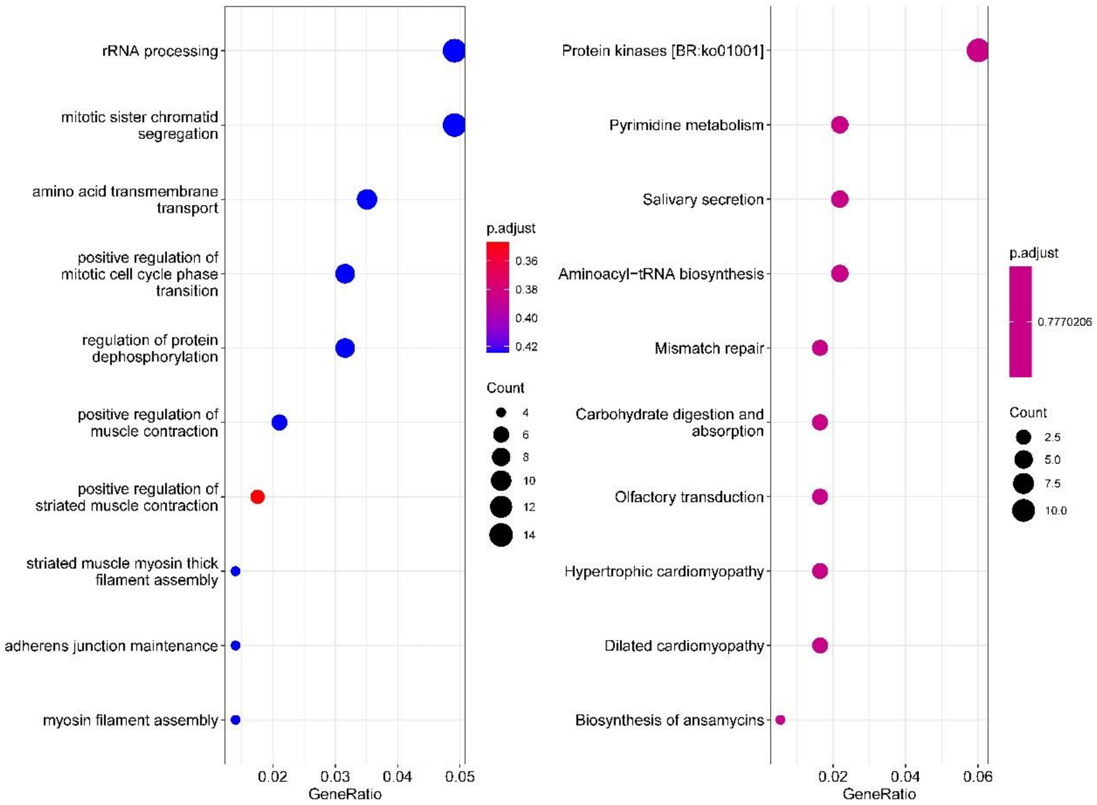


**Fig S6. Function enrichment of positive selected genes.** Left is GO terms; right is KEGG terms.

**Table S1. Gene number, gene density and intron length of each chromosome.**

| chr | length | gene number | gene density | mean intron length |
|---|---|---|---|---|
| LG1 | 991018619 | 3157 | 3.19E-06 | 15383 |
| LG2 | 901651818 | 3040 | 3.37E-06 | 15174 |
| LG3/X | 806237231 | 1985 | 2.46E-06 | 18258 |
| LG4 | 689659731 | 2247 | 3.26E-06 | 15184 |
| LG5 | 581103692 | 1850 | 3.18E-06 | 15713 |
| LG6 | 485803872 | 1439 | 2.96E-06 | 16076 |
| LG7 | 427868946 | 1218 | 2.85E-06 | 16448 |
| LG8 | 418016564 | 1421 | 3.40E-06 | 15127 |
| LG9 | 377406850 | 2047 | 5.42E-06 | 15032 |
| LG10 | 148047574 | 732 | 4.94E-06 | 16256 |
| LG11 | 142112905 | 638 | 4.49E-06 | 13794 |
| LG12 | 134840001 | 493 | 3.66E-06 | 12899 |

**Table S2. $d_N$, $d_S$ and $d_N/d_S$ values of different categories.**

| Category | $d_N$ | $d_S$ | $d_N/d_S$ |
|---|---|---|---|
| X-conserved | 0.015714[d] | 0.137885[d] | 0.101397[e] |
| A-X | 0.028111[c] | 0.190683[c] | 0.144579[d] |
| X-X | 0.028380[c] | 0.166257[d] | 0.159734[c] |
| A-A | 0.041803[b] | 0.213579[b] | 0.190432[b] |
| X-Lmig | 0.103788[a] | 0.290710[a] | 0.372944[a] |
